# Branch extension and pruning share the same regulatory module in the developing *Drosophila* airways

**DOI:** 10.64898/2026.04.24.720541

**Authors:** Ryo Matsuda, Chie Hosono, Kaoru Saigo, Christos Samakovlis

## Abstract

Branched tubular organs represent a common solution to the problem of fluid transport in large animals. The growth of new branches has been extensively studied but the regulation of branch pruning remains underexplored. Here, we investigate branch removal in the stereotyped branching patterns of the developing *Drosophila* airways. After progenitor invagination, the tips of the distal airways generate a stereotyped branching pattern in each central metameric unit. An intriguing exception in the repeating patterns is the lack of branches targeting the visceral mesoderm (visceral primary branch, VB) in the third and nineth metameres. We show that these branches initially form but the localized expression of the pro-apoptotic gene *reaper* and resultant apoptosis prune them. We reveal that VB3/9 pruning entails four sequential programs. First, a common distal outgrowth program promotes budding and extension of the primary branches, including VB3 and 9. Second, a VB identity program is established representing the ground state of all primary branches. Third, the Bithorax-Complex transcription factors define metameric identities and interfere with the VB program to induce *reaper* and apoptosis specifically in VB3/9 in a concentration dependent manner. Finally, the default VB cell identity is transformed by extrinsic BMP/Decapentaplegic and WNT/Wingless into more derived primary branch identities and spared from pruning in metamers 3 and 9. Our results demonstrate that molecular and genetic circuits promoting branch emergence and extension can be regionally modified and deployed also for branch pruning.

**Author summary:** Many of our internal organs like the lung, kidney and various glands are composed of epithelial tubes. The function of these organs is sustained by the morphologies of the constituting tubes. The formation of distinct branching patterns in tubular networks including regional variegations can be accomplished by branch ramification and branch pruning. Extensive studies of tube ramification and extension showed that they are typically regulated by extrinsic guidance cues. In contrast to branch extension, branch maintenance and removal is much less understood. Here, we identify a program that intrinsically promotes apoptosis to prune specific branches in the developing *Drosophila* airways. Intriguingly, the branch extension program is diverted to a branch pruning program. Activation of the branch outgrowth program is prerequisite for both branch extension and pruning. The Hox gene expression code that characterizes regional cell identities along the body axes intrinsically interferes with the branch extension program of visceral primary branches (VB) targeting the visceral muscles. This results in VB pruning only in metameres 3 and 9, where extrinsic WNT and BMP guidance cues direct towards distal branch identities that are spared from apoptotic pruning. Similar interplay of branching and pruning programs may operate in sculpting our lung and vasculature.

## Introduction

Humans and *Drosophila* fruit flies possess similar respiratory systems, although vertebrates and insects are very divergent animal groups. Both systems are dominantly composed of epithelial tube networks (Romero et al. 2025). In humans, oxygen flows through the airway tubes to the distal tips of lung ramifications called alveoli (Herriges and Morrisey 2014; Kishimoto and Morimoto 2021). There, oxygen is transferred to erythrocytes in specialized capillaries and becomes transported through the extensively ramified vascular network to the whole body (Potente and Makinen 2017). In *Drosophila*, on the other hand, a highly ramified epithelial tube network called the tracheal system enables airflow to the target cells inside the whole body (Manning and Krasnow 1993; Samakovlis et al. 1996a; Samakovlis et al. 1996b; Hu and Castelli-Gair 1999; Ghabrial et al. 2003). Blood cells in the open hemolymph transiently attach to the fine capillaries at the distal airway tips and assist oxygen delivery to target cells (Shin et al. 2024).

The cellular mechanisms promoting tube generation and branching include sequential events of new sprout formation followed by maintenance or removal of nascent branches. The former process has been studied extensively, leading to a generalized view that extracellular guidance molecules attract or repel growth of tubes by signaling through their respective receptors expressed in the cells of the network (Ghabrial et al. 2003; Affolter and Caussinus 2008; Eilken and Adams 2010; Romero et al. 2025).

In contrast, the genetic and molecular analysis of branch maintenance and pruning has been less investigated (Levi et al. 2006; Chen et al. 2012; Korn et al. 2014; Franco et al. 2015; Lenard et al. 2015; Zhang et al. 2018). Whereas flow-dependent and -independent pruning operate during angiogenesis, the developing *Drosophila* airway tubes are presumed to be devoid of shear stress.

The *Drosophila* airways originate from 10 metameric primordial cell clusters specified at left and right sides of the embryonic ectoderm (Perrimon et al. 1991; Manning and Krasnow 1993; Samakovlis et al. 1996a; Hu and Castelli-Gair 1999). Radially confined sequential invagination of the progenitors in each 2-dimensional primordium generates a primitive cavity harboring proximal and distal regional differences (Brodu and Casanova 2006; Nishimura et al. 2007; Matsuda et al. 2015b; Matsuda et al. 2026). The tips of the distal airways typically extend 6 primary branches and some of them fuse with the contralateral or ipsilateral neighbors to interconnect the system (Figure 1A) (Perrimon et al. 1991; Manning and Krasnow 1993; Samakovlis et al. 1996a; Samakovlis et al. 1996b). In spite of the stereotyped primary branching of each metamere, branches targeting the visceral muscles (VBs) are missing in metamere 3, 9 and 10 (Manning and Krasnow 1993). This pattern deviation is investigated in this report.

**Figure 1.**
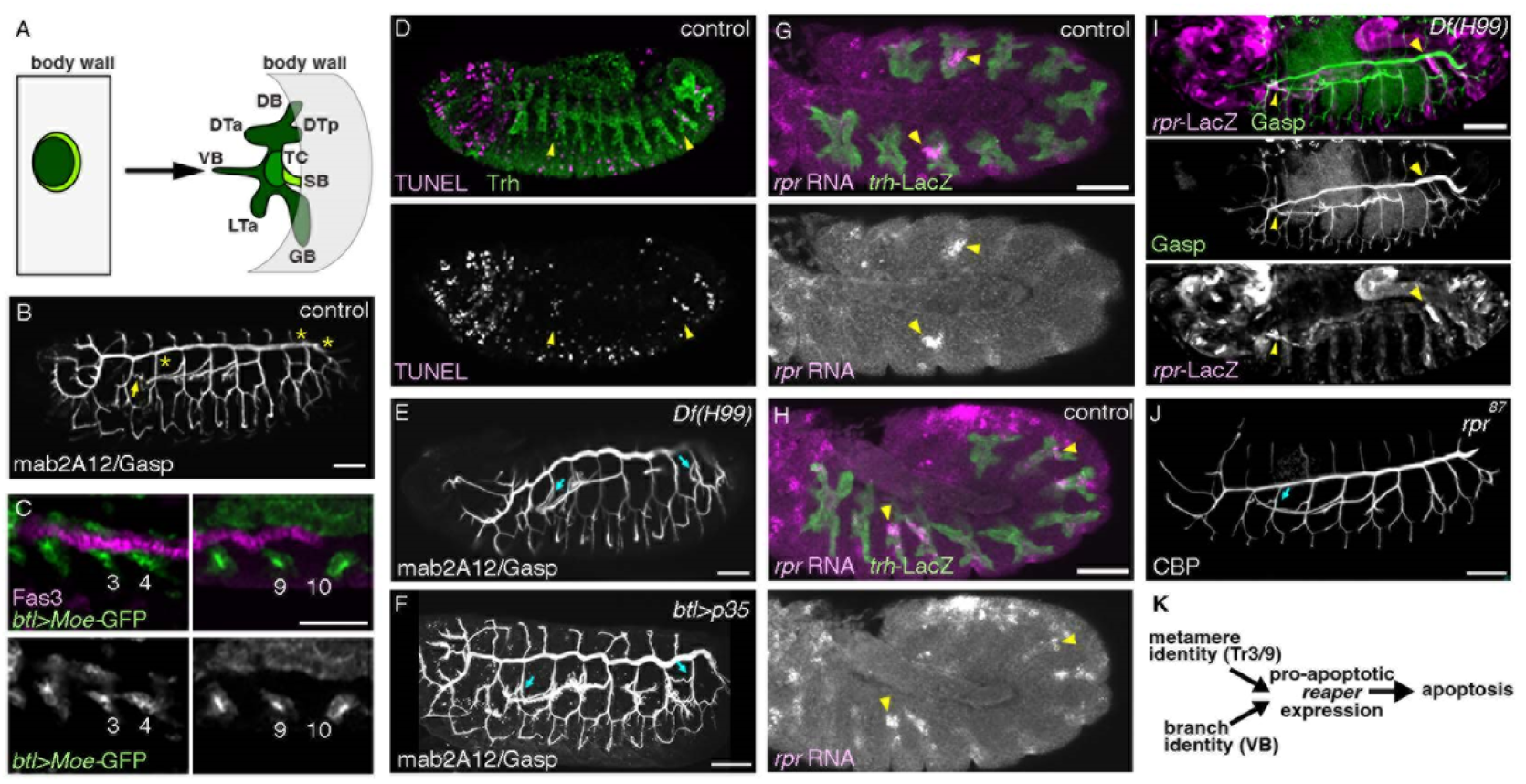
Apoptotic pruning of VB3 and VB9. (A-C) The developmental origin of the missing VB3/9/10. Primordium invagination and their primary branching pattern in a typical metamere is shown in A. DB (dorsal branch), DTa (dorsal trunk anterior), DTp (dorsal trunk posterior), VB (visceral branch), LTa (lateral trunk anterior), GB (ganglionic branch), TC (transverse connectives), SB (spiracular branch). A projection of a stage 16 embryo stained with mab2A12 (B) showing missing VB3/9/10 (asterisks), compared to a typical VB (arrows). At stages 11/12, VB precursors marked with *btl>moe-GFP* extending to Fas3 positive visceral mesoderm are detected in Tr3 and Tr9, but not in VB10 (C). (D-K) *rpr* mediated apoptotic pruning of VB3/9. TUNEL signals (D) are evident at VB3/9 during stages 12-13 (arrowheads). Suppression of apoptosis by *H99* deficiency (E) or by overexpression of *p35* (F) restores VB3/9 formation (arrows). *rpr* transcripts in the control (G-H) or *rpr-lacZ* reporter activity in *H99* mutants (I) are detected preferentially in VB3/9 (arrowheads). *rpr* mutants variably restores VB3/9 formation. A working model of VB3/9 pruning regulation (K). Scale bars, 50μm.

Two types of signaling systems control primary branching and branch identities. The first type is required for all the primary branches and the second regulates distinct primary branches. Distalizing factors including *hedgehog*, *ventral veinless (vvl)*/*drifter*/*u-turn*, *rhomboid* (*rho*) and *breathless* (*btl*)/*FGFR* promote airway tube formation and its ramification (Matsuda et al. 2015b; Matsuda et al. 2026). For example, Branchless (Bnl)/FGF released from surrounding tissues guides directional extension of all the primary branches by activating Breathless (Btl)/FGFR expressed on the airway cells (Glazer and Shilo 1991; Klambt et al. 1992; Sutherland et al. 1996). In parallel, a second regionally restricted system composed of WNT and BMP members induce and guide distinct primary branch identities. Wingless (Wg) and Wnt2 expressed in transverse stripes of the overlying ectoderm upregulate the expression of *spalt-major* (*salm*), encoding a zinc-finger transcription factor (TF) and specify the short and thick nature of the dorsal trunk (DT) (Kuhnlein and Schuh 1996; Chihara and Hayashi 2000; Llimargas 2000; Llimargas and Lawrence 2001; Ribeiro et al. 2004; Shaye et al. 2008; Matsuda et al. 2015a). In contrast, Decapentaplegic (Dpp)/BMP expressed in dorsal and lateral longitudinal ectodermal stripes induces expression of the *knirps* (*kni*) and *knirps-related* (*knrl*) TFs (Vincent et al. 1997; Chen et al. 1998; Ribeiro et al. 2002). This signaling module selects the dorsally migrating branch (dorsal branch/DB) and specifies the ventrally migrating branches (lateral trunk/LT and ganglionic branch/GB). The visceral branch (VB) targets the visceral muscles and co-expresses the TFs *empty spiracle* (*ems*) (Ebner et al. 2002) and *kni/knrl*, the latter of which is induced independently of Dpp/BMP signaling in the VBs (Vincent et al. 1997). Previous studies found that ectopic activation of Wg/WNT signaling transforms VB into DT (Chihara and Hayashi 2000; Llimargas 2000). However, the endogenous mechanisms for selecting VBs remain unexplored.

Hox genes encode an evolutionally conserved family of TFs that confers distinct morphological identities to organs along the entire body (Lewis 1978; Kaufman et al. 1980; Pearson et al. 2005; Maeda and Karch 2006), whose regulatory targets include pro-apoptotic genes (Lohmann et al. 2002; Domsch et al. 2015). Hox genes can directly activate or repress some targets whereas Hox DNA-binding specificity for other targets is aided by a pair of cofactors, *extradenticle* (*exd*) and *homothorax* (*hth*) or various interacting TFs (Mann et al. 2009; Sorge et al. 2012; Merabet and Mann 2016; Bischof et al. 2018). The *Drosophila* genome contains two clusters of Hox genes, the Antennapedia complex and the Bithorax complex (BX-C) (Lewis 1978; Kaufman et al. 1980). The ectoderm and its derivatives, appendages and the airway primordia (Carroll 1995; Sanchez-Higueras et al. 2014; Matsuda et al. 2015a; Barcenilla-Merino et al. 2025) express Hox genes in register with para-segments (Maeda and Karch 2006), developmental units along the AP axis of the *Drosophila* body (Martinez-Arias and Lawrence 1985). In the context of this study, we focused in BX-C genes, whose 3 products (*Ultrabithorax*/*Ubx*, *abdominal A*/*abdA* and *Abdmoinal B*/*AbdB*) are differentially expressed in tracheal metameres 2-10 (Tr2-10) (Matsuda et al. 2015a).

We identified apoptotic pruning as a morphogenetic mechanism shaping the branching diversities of the *Drosophila* airways. We present a genetic circuit where 3 programs of primordial cell invagination, primary branch diversification and metamere identities orchestrate developmental pruning of VB3/9. The acquisition of primary branch identity is a prerequisite for pruning of those branches. We propose that the actions of distalizing factors promoting primary branch extension are modified by regional identity programs to induce subsequent localized apoptotic pruning.

## Results and Discussion

### VB3/9 is pruned by apoptosis

*Drosophila* airway branching has a standard design for all central metameres, except for the VB3 and VB9 and additionally VB10, which are not detectable at stage 16, after major airway branching patterns are established (Figure 1A,B) (Manning and Krasnow 1993). To identify the underlying mechanisms of these missing VB3/9/10, we followed the branching of the Tr3/9/10 metameres at different embryonic stages. At late stage 11, when primary branching has initiated VB3 touches its target tissue, the visceral mesoderm, and extends anteriorly on it (Figure 1C). VB9 sprouting straight toward its target also occurs but is more difficult to detect (Figure 1C). We were unable to detect VB10 formation (Figure 1C). By early stage 13, scattered dots of cytoplasmic markers of the airway cells become detected at locations of VB3/9 (Figure S1), suggesting apoptotic cell degradation of VB3/9. Consistently, by late stage 12, we detected localized DNA fragmentation signals by TUNEL in VB3/9 (Figure 1D). These results suggest that VB3/9 are initially formed but lost due to apoptotic pruning. In contrast, the lack of VB10 originates from failure in specifying the VB10 precursors.

To test the potential roles of apoptosis in VB3/9 loss, we examined mutants of *H99* deficiency, uncovering 3 major pro-apoptotic genes, *rpr (White et al. 1994)*, *grim* (Chen et al. 1996) and *head involution defective* (Grether et al. 1995), thereby suppressing all embryonic developmental apoptosis (White et al. 1994). In *H99* mutants, both VB3/9 are restored, while VB10 is not (Figure 1E). Consistently, airway-specific overexpression of *p35*, a baculovirus inhibitor of apoptotic caspases (Hay et al. 1994) variably restores VB3/9 (Figure 1F). We conclude that VB3/9 initially form, but are pruned through apoptosis induction by stage 13.

### *rpr* expression in VB3/9 foresees their pruning

Out of the 3 pro-apoptotic genes uncovered by the *H99* chromosomal deficiency, we noticed that *rpr* transcripts are specifically induced in VB3/9 at early stage 12 (Figure 1G). Slightly later, *rpr* RNA becomes variably detected also in VB4 and VB8 (Figure 1H).

To further investigate the transcriptional *rpr* activation in the airways we used a *rpr-lacZ* reporter construct (Nordstrom et al. 1996) and analyzed its expression in the *H99* mutant background where apoptotic pruning of VB3/9 is suppressed. Expression of *rpr-11kb-lacZ* is persistently detected in the restored VB3/9 of H99 mutants (Figure 1I). VB3/9 are partially restored in *rpr* mutants (Figure 1J) suggesting the functional role of *rpr* in shaping VB3/9. The partial rescue of VB3/9 in *rpr* mutants may reflect the additional involvement of other pro-apoptotic genes that act redundantly in VB pruning.

The induction of *rpr* expression specifically in VBs of only selected metameres suggest that there are two transcriptional inputs controlling apoptotic pruning (Figure 1K). One input deriving from the metamere-identity genes and the other from the branch-identity gene hierarchy.

### The three routes of primary branching are exposed to different combinations of Wg/Wnt2 and Dpp/BMP guidance signals

To characterize the branch identity hierarchy for VB3/9 pruning, we first clarify how the airway progenitors diversify their identities during the initial phases of differentiation. It is deduced that the cells in the central parts of the 2D primordia would constitute the tips of the distal airways, giving rise to the primary branches (Figure 2A) (Matsuda et al. 2015b). The primary branch progenitors are divided into 3 types, based on *hairy* expression in the dorsal-central parts of the 2D primordia and *Drop* (*Dr*)*/muscle specific homeobox* (*msh*) expression in the ventro-lateral ectoderm (Figure 2A) (Matsuda et al. 2015b; Matsuda et al. 2026). The progenitor cells expressing *hairy* but not *Dr/msh* (*hairy^+^ Dr/msh^-^* progenitors) generate DBs and DTs whereas *hairy^+^ Dr/msh^+^* double positive progenitors generate VBs. *hairy^-^ Dr/msh^+^* progenitors would generate the ventrally migrating primary branches.

**Figure 2.**
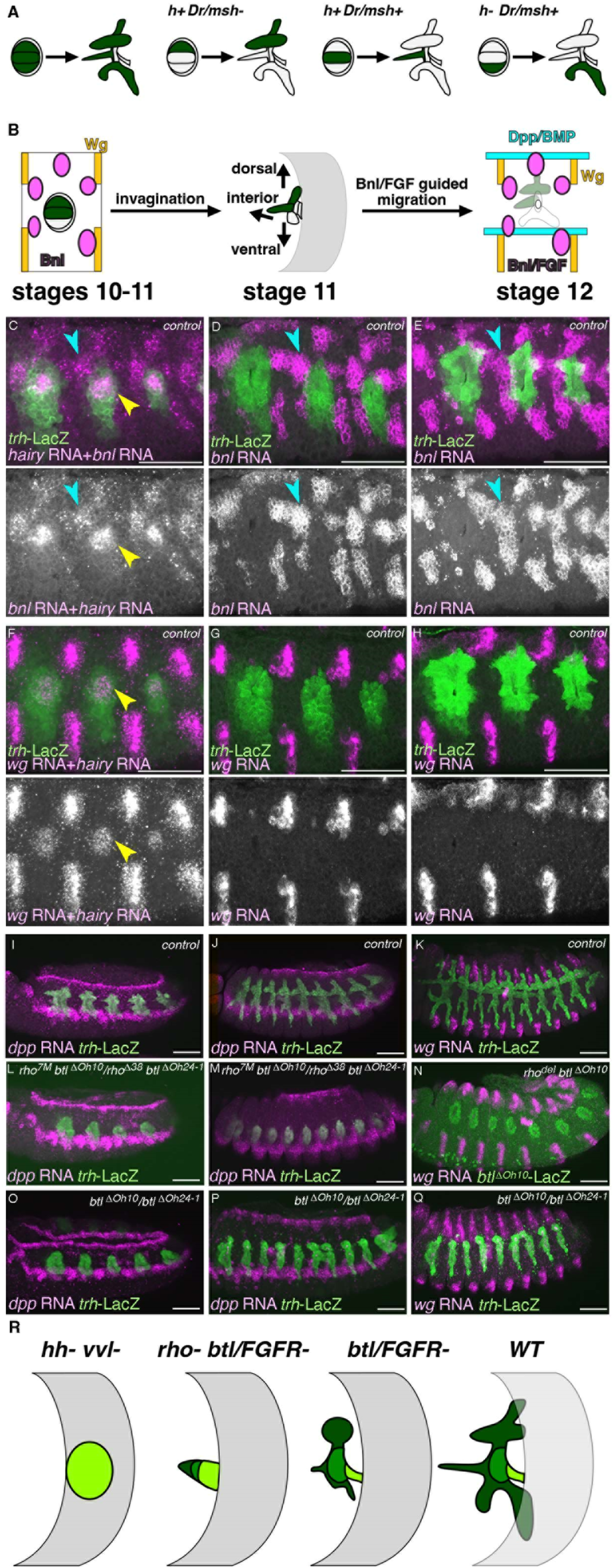
Three migratory routes of the airway progenitors along different combinations of guidance molecules and their regulation by EGFR and btl/FGFR. (A) A conceptual fate map of the 2D airway primordia. Expression of *hairy* and *Dr/msh* along the dorso-ventral axis subdivides the distal progenitors into 3 kinds, DB/DT, VB and LT/GB, corresponding to dorsal, interior and ventral directions of migration. (B-K) Locations of the airway progenitors relative to the expression domains of the 3 guidance molecules, *bnl/FGF*, *wg/Wnt* and *dpp/BMP*, before and after invagination are summarized in (B). Before invagination, the *hairy* expressing DB/DT/VB progenitors in the 2D primordia are located ventral to the expression domains of *bnl/FGF* and *wg/Wnt* that guide future DT migration (B left, stages 10-11). Invagination transforms the dorso-ventral patterning of the progenitors into three routes of branch extension, dorsal, interior and ventral (B middle, stage 11). Dorsal-ward migration of the invaginated DB/DT progenitors initially exposes them Bnl/FGF and Wg/Wnt. Then, some of them encounter Dpp/BMP (B right, stage 12). On the other hand, the invaginated progenitors of the ventrally migrating branches (LT, GB) initially encounter Bnl/FGF and Dpp/BMP. Then, some of them encounter Wg/Wnt. The invaginated progenitors of the interiorly migrating branches (VB) encounter Bnl/FGF. Expression of *bnl* (C-E), *wg* (F-H, K) or *dpp* (I-J) in the control is shown. Note that the airway progenitors are before invagination in (C, F) and that embryos are in progressively later stages in (D, G), (E, H), (I, J) and K. The *hairy* expressing DB/DT/VB progenitors are marked by yellow arrowheads in (C, F) whereas the DTa-guiding *bnl* expression domains are marked with blue arrowheads in (C-E). (L-R) Regulation of progenitor location by RTKs. Expression of *dpp* (L, M, O, P) or *wg* (N, Q) is used as reference locations. The airway progenitors stay at the invaginated places in *rho btl/FGFR* double mutants by stage 13 (L-N, R). Comparing *btl/FGFR* mutants (O-Q, R) with *rho btl/FGFR* double mutants, *btl/FGFR* mutant cells migrate in all 3 directions (dorsal, interior or ventral), corresponding to their apparent structures of DT, VB and GB. Scale bars, 50μm.

Double labeling of *hairy* and *bnl* or *wg/WNT* transcripts show that the DB/DT progenitors are located ventrally to the *bnl* expression that guides the DT cells (Figure 2B, C, F). The invaginated DB/DT progenitors, which move dorsally under the overlying ectoderm are presumed to receive Bnl/FGF and WNTs (Wg and Wnt2) secreted from the overlying ectoderm, the latter inducing the DT identity (Figure 2B, D-E, G-H) (Sutherland et al. 1996; Chihara and Hayashi 2000; Llimargas 2000; Llimargas and Lawrence 2001). Then, the dorsally located cells of these DB/DT progenitors become exposed to Dpp/BMP secreted from the dorsal part of the embryos (Figure 2B, I-J) (Vincent et al. 1997; Chen et al. 1998). On the other hand, the *hairy ^-^Dr/msh^+^* progenitors of ventrally migrating branches receive Dpp/BMP, Bnl/FGF and Wg/WNT from the overlying ectoderm (Figure 2B, D-E, G-H, I-K) whereas the internally migrating *hairy^+^ Dr/msh^+^* VB progenitors are only exposed to Bnl/FGF (Figure 2B).

In the absence of the two distalizing factors Hh and Vvl, many of the airway progenitors fail to invaginate (Matsuda et al. 2026) and they also fail to express Kni as early as at stage 11 (Figure S2A-B). On the other hand, in the absence of another pair of the distalizing factors *rho* and *btl/FGFR*, the invaginated cells stay at the original places by stage 13, which is presumably out of the effective range of the DT inducing Wg/Wnt2 signals (Figure 2N, R) but are surrounded by the ventral stripe of *dpp/BMP* expression (Figure 2L, M, R). Still, Kni expression is variably affected (Figure S2C).

In *btl/FGFR* mutant, where Bnl/FGF guided progenitor migration does not occur, 3 remnant structures of the primary branches are obvious in the tips of the distal airways, DT, VB and GB, which correspond to the 3 subdivision of the central parts of the 2D primordia along the dorso-ventral axis of the embrtyos, *hairy^+^ Dr/msh^-^*, *hairy^+^ Dr/msh^+^* and *hairy^-^Dr/msh^+^* progenitors (Figure 2A, O-R). Though DB is not structurally apparent in *btl/FGFR* mutants, *kni* as well as *kni-(dpp)-lacZ*, a *kni* enhancer reporter that responds to Dpp/BMP are variably induced in the dorsal tips of the DT as well as in GB (Figure S2D) (Matsuda et al. 2015a).

Taken together, whereas invagination transforms the centro-peripheral patterning of the 2D primordia into the proximo-distal differences of the 3D tubes (Matsuda et al. 2015b; Matsuda et al. 2026), the dorso-ventral patterning of the progenitor field would be converted into the 3 prominent directions of vessel extension, dorsal, interior and ventral. Extrinsic signals including Wg/Wnt2, Dpp/BMP and Bnl/FGF collaborate to diversify the direction of the distal airway tips to promote primary branching and establish the different branch identities (Figure 2A, B, R).

### VB3/9 branches are restored by overexpression of the VB selector gene *ems*

To identify the anticipated visceral branch identity inputs to VB3/9 pruning, we focused on the transcription factors *kni/knrl* and *ems*, which are co-expressed in VBs (Figure 3A, B) (Vincent et al. 1997; Chen et al. 1998; Ebner et al. 2002). We first examined effects of overexpressing Kni or Ems on VB growth and maintenance. Overexpression of Kni and Knrl with *btl-*Gal4 caused loss or thinning of the DT branches, as reported previously (Figure 3C) (Chen et al. 1998) but had no effects on the VB3/9 pruning. Notably however, overexpression of Ems in the airways generated VB-like extensions not only in Tr3/9 but also in Tr10 (Figure 3D) suggesting an essential role of *ems* in VB specification and pruning. To examine if *ems* overexpression impacts on induction of *rpr* in VB3/9, we monitored *rpr* expression in *H99* mutant background. The strong *rpr-lacZ* expression detected in VB3/9 of the control embryos (Figure 1I) or upon *kni* overexpression (Figure 3E) was abolished by Ems overexpression (Figure 3F). This suggests that Ems overexpression restores VB3/9 by inhibiting *rpr* induction and subsequent apoptosis.

**Figure 3.**
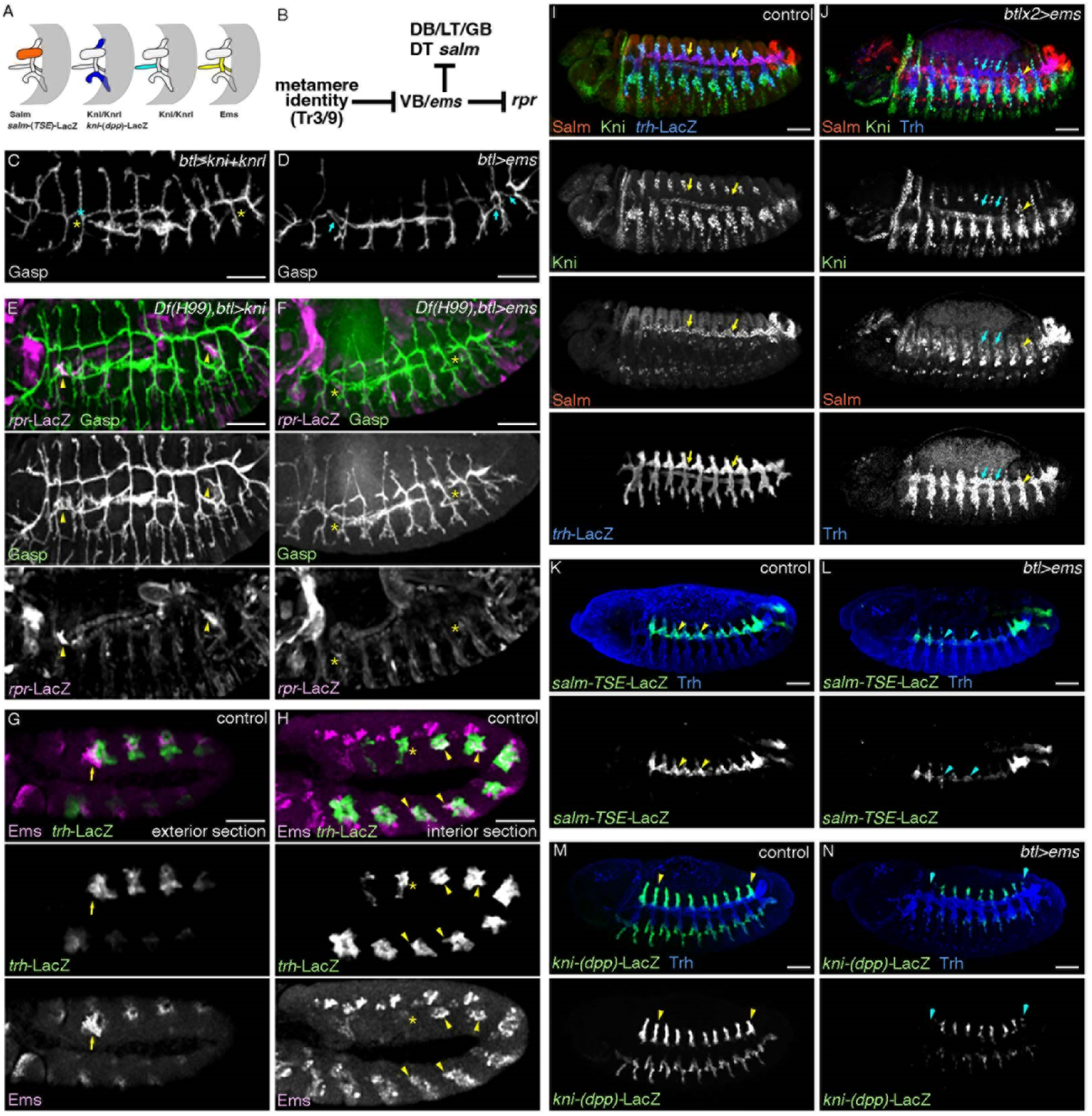
The VB selector gene *ems* is involved in VB loss. (A-F) *ems* overexpression causes VB3/9/10 formation. Differential expression of *salm*, *kni/knrl* and *ems* highlight distinct primary branches (A). A regulatory model of VB3/9 pruning by the VB selector gene *ems* (B). The airway specific overexpression of Kni and Knrl (C) does not restore the missing VB3/9/10 (asterisks). Upon Ems overexpression (D), all of VB3/9/10 are formed (arrows). In Df(*H99)* background, compared to Kni overexpression (E, arrowheads), Ems overexpression abolishes *rpr-lacZ* expression in VB3/9 (F, asterisks). (G-H) Ems expression. In the wild type at stage 11, a new Ems expression domain in the ectoderm appears in the posterior neighbor of Tr10 (G, arrows). More interiorly (H), budding of VB10 is not detectable while Ems expression in the short VB9 bud is very low (asterisks). In the more anterior metameres including VB3, Ems expression in VB is comparable (arrowheads). (I-N) Effects of Ems overexpression on the primary branch identities. Compared to the control (I, K and M), Ems overexpression abolishes or reduces expression of Salm (J, arrows), *salm-TSE-lacZ* (L, arrowheads) or *kni-dpp-lacZ* (N, arrowheads), with occasional ectopic induction of Kni in the DT regions (J, arrowheads). Scale bars, 50μm.

To further test how *ems* affects VB3/9 pruning, we studied Ems expression in VBs. At early stage 11, Ems is expressed broadly in the main airway primordia (Dalton et al. 1989; Walldorf and Gehring 1992), and by stage 13 its expression is confined to VB, TC (Ebner et al. 2002) and SB in a typical central metamere (Figure S3A). At late stage 11 when Ems expression resolved, the central/distal parts lose Ems expression and its expression initiates in VBs. At this stage, Tr9/10 showed a clear deviation. In Tr10, an expression domain corresponding to a part of the posterior spiracle primordium became evident in the ectoderm (Figure 3G) (Dalton et al. 1989; Walldorf and Gehring 1992; Jones and McGinnis 1993). In the distal part of Tr10, it was hard to detect a VB-like extension and Ems expression (Figure 3G, H). In Tr9 on the other hand, a small VB-like bud was detectable but showed very low Ems expression (Figure 3H). In contrast, Ems expression was largely comparable in all other VBs including VB3 (Figure 3H). These results suggest that reduction or loss of Ems in the distal parts of Tr9 and Tr10 is responsible for VB9 pruning and VB10 absence. VB3 pruning may involve posttranslational mechanisms interfering with *<u>ems</u>*functions.

In addition to the ectopic VB3/9/10, Ems overexpression caused stalling or loss of other primary branches (Figure 3D, F). *salm* and *salm-TSE-lacZ*, are two markers for DT identity (Kuhnlein and Schuh 1996; Chen et al. 1998) whereas *kni-(dpp)-lacZ* is a marker for the DB, LT and GB cells (Chen et al. 1998). Ems overexpression reduced or abolished expression of *salm, salm-TSE-lacZ and kni-(dpp)-lacZ by* stage 13 (Figure 3 I-N). In contrast, Kni protein expression remained comparable to the control in the stalled DB/ LT/GB regions (Figure 3I, J) and it occasionally expanded to the DT region at stage 13 (Figure 3J).

Moreover, the fusion cell identity monitored by expression of Dys, a TF regulating branch fusion (Jiang and Crews 2003) was variably lost upon Ems overexpression (Figure S3B-C). The terminal cell identity, monitored by DSRF expression (Affolter et al. 1994a; Guillemin et al. 1996) was mildly affected under this condition (Figure S3B-C). These results show that overexpressed Ems represses the Wg/WNT and the Dpp/BMP derived primary branch identities and promotes the VB identity, where branch tips are devoid of fusion cells. Persistent Kni expression in the presence of overexpressed Ems proteins would be realized by a VB-specific enhancer activity, while the Dpp responsive airway enhancers of *kni* are repressed by Ems. In contrast to the function of overexpressed Ems to promote the VB identity, an *ems* loss of function mutation only variably allows VB formation (Ebner et al. 2002). We speculate that like *kni/knrl* paralogues, *ems* functions could be compensated by *E5*, an Ems paralogue that is expressed in the airway progenitors in a way similar to *ems* (Dalton et al. 1989; Hammonds et al. 2013).

Taken together, the abilities of missexpressed Ems to induce ectopic VBs and to suppress various characteristics of the other primary branch identities suggest that *ems* is a selector gene for VB. We propose that VB3/9 pruning reflects interference with the VB identity in these metameres.

### Wg/WNT and Dpp/BMP signaling protects other primary branches from pruning in Tr3/9

The localized expression of *rpr-lacZ* only in the VB and not in other primary branches in Tr3 and 9 prompted us to ask how the remaining primary branches are protected from apoptosis in these metameres. Previous studies have revealed that extrinsic signals of Wg/WNT and Dpp/BMP variably affect VB formation (Vincent et al. 1997; Chihara and Hayashi 2000; Llimargas 2000). We therefore monitored the VB cell markers, Ems and Kni, in loss- and gain-of-function mutants affecting Dpp/BMP and Wg/WNT signaling components.

Upon Wg over expression by *btl-*Gal4, the expression domains of both Kni (Llimargas 2000) and Ems became reduced (Figure 4 A-B). This was accompanied by an increase of Salm-positive DT cells (Chihara and Hayashi 2000; Llimargas 2000), corroborating the notion that ectopic Wg/WNT signaling transforms VB into DT. Conversely, in mutants of *armadillo* (*arm*), a mediator of Wg/WNT signaling (Noordermeer et al. 1994; Siegfried et al. 1994), Kni expression was expanded to the DT region (Figure 4C) (Llimargas 2000). The ectopic Kni induction was not accompanied by expanded expression of *kni-(dpp)-lacZ* (Figure 4D). This suggests that the Dpp/BMP signaling range at the DT region is largely intact in *arm* mutants. Like Kni, Ems expression was significantly expanded to cover the DT region in *arm* mutants (Figure 4D). These results suggest that the DT cells are transformed to the VB identity upon reduction of Wg/WNT signaling. We conclude that Wg/WNT signaling, also at endogenous levels transforms the VB cells into the DT cell type.

**Figure 4.**
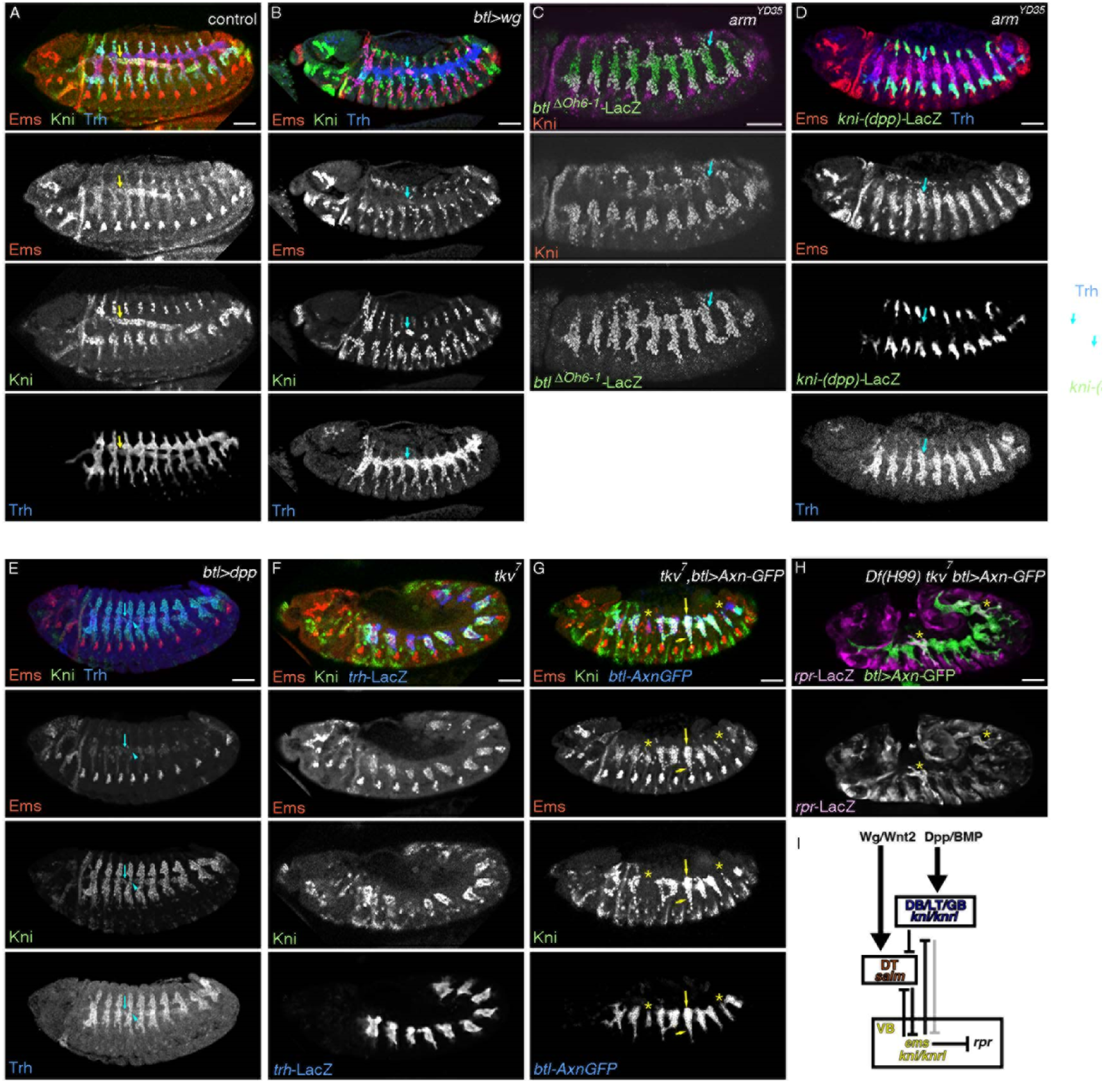
Regulation of VB identity and VB pruning by Wg/WNT and Dpp/BMP signaling. Compared to the control (A), the airway specific overexpression of Wg (B) reduces expression of both Kni and Ems (arrows). In *arm* mutant (C, D), both Kni (C) and Ems (D) are induced at the DT position (arrows) without changing the distribution of *kni-(dpp)-*LacZ (D). Dpp/BMP overexpression (E) reduces Ems expression (arrows) whereas Kni is ectopically induced as reported. Compared to tkv mutants (F), expression of Axn-GFP in *tkv* mutants (G) causes expansion of both Kni and Ems to most primary branches and prominent reduction of Trh+ cell number in Tr3/9. This accompanies expansion of *rpr-lacZ* to all distal parts of Tr3/9 in *Df(H99)* mutant background (I). Asterisks in (G-H) mark VB3/9. Regulatory interactions among the Wg/Wnt2-Salm module, the Dpp/BMP-Kni/Knrl module and the VB state are shown in (I). Scale bars, 50 μm.

Wg/WNT signaling upregulates *salm* expression in the airway progenitors (Chihara and Hayashi 2000; Llimargas 2000). We therefore tested if *salm* has the same effects on Ems and Kni expression as *wg*/*WNT* signaling. Overexpression of Salm with *btl-*Gal4 reduced the expression of both Ems and Kni in the VB regions (Figure S4A). Unexpectedly however, *kni-(dpp)-lacZ* expression was expanded in *salm* mutants, suggesting that the Dpp/BMP signaling range was correspondingly expanded (Figure S4B,C). Indeed, ectopic *dpp/BMP* expression was detected as transverse stripes in the lateral ectoderm of *salm* mutants (Figure S4D-H) from stage 11 on, suggesting that *salm* expression in the dorsal ectoderm (Kuhnlein and Schuh 1996) functions to repress ectopic *dpp/BMP* expression in the lateral ectoderm. Collectively, the Salm overexpression phenotypes support the notion that Wg/WNT signaling and the resultant Salm upregulation in the DT cells operate to repress the VB state.

On the other hand, endogenous Dpp induces *kni* expression in the primary branches migrating dorsally and ventrally (DB, LT, GB) through Dpp-responsive enhancers (Vincent et al. 1997; Chen et al. 1998). Dpp overexpression induces *kni* in all the distal cells (Vincent et al. 1997) whereas ectopic *kni-(dpp)-lacZ* expression was also detected in VBs (Figure S4I). Furthermore, Ems expression in VB was variably reduced (Figure 4E). This argues that that Dpp/BMP signaling can interfere with the VB identity. We further examined the effects of loss of *dpp/BMP* signaling. In mutants for *thick veins (tkv)*, one of the heterodimeric Dpp/BMP receptors (Affolter et al. 1994b; Brummel et al. 1994; Penton et al. 1994), primary branch formation showed a variety of defects (Affolter et al. 1994b). DB formation was largely suppressed in *tkv* mutants as *tkv* controls both *kni* induction in the DB (Vincent et al. 1997; Ribeiro et al. 2002) and *bnl* expression attracting DB migration (Vincent et al. 1997). Similarly, in the ventral branches of *tkv* mutants, *kni* expression was variably lost (Vincent et al. 1997) and concomitantly, *salm* expression was often detected ectopically in cells near fusion points of the LT, close to the source of ectodermal Wg expression (Figure S4J). This suggests that the endogenous Dpp/BMP signaling antagonizes Wg/WNT signaling, not only in DB (Chen et al. 1998) but also in the ventral part of the airways, as proposed originally by Llimargas (Llimargas 2000; Matsuda et al. 2015a).

While the extension of the ventral primary branches is defective in *tkv* mutants (Vincent et al. 1997), we noticed that Ems and Kni were variably extended also in the remaining ventral primary branches (Figure 4F, S4K). These results suggest that Dpp/BMP signaling restricts expression of Ems and the VB state.

*kni* and *knrl* are transcriptional targets and mediators of Dpp/BMP signaling but also define the VB identity. Removal of both *kni* and *knrl*, in *Df(kni/knrl)* mutants, variably affected VB formation causing a range from VB loss to stalling phenotypes (Chen et al. 1998), suggesting that *kni/knrl* promote VB specification. However, the residual VB branches in *Df(kni/knrl)* mutants, still retained Ems expression (Figure S4L), suggesting a redundant mode in VB establishment. In addition, we detected variable ectopic expression of Ems in LT/GB cells of *Df(kni/knrl)* mutants whose segmentation defects are rescued by a transgene (Figure S4M-N). This may reflect their role in *ems* repression downstream of Dpp/BMP.

Finally, in *arm*;*tkv* double mutants, where both Wg/WNT and Dpp/BMP signaling are reduced, VBs are the only primary branches that emanate from the TC (Llimargas 2000). To further examine the identities of the airway cells upon reductions of Wg/WNT and Dpp/BMP signaling, we overexpressed in *tkv* mutants a GFP fused variant of Axin (Axn), a negative regulator of Wg/WNT signaling (Hamada et al. 1999; Willert et al. 1999; Cliffe et al. 2003). Under this condition, we detected 3 types of cell populations in a typical metamere. The proximal part of the airways or SBs (spiracular branch) were occupied with Ems^+^GFP^-^ cells. More distally located were Ems^+^GFP^+^ cells corresponding to TC (transverse connective). In the distal part of the airways, most Axn-GFP positive cells are also positive for Ems and Kni (Figure 4G). This indicates that all the primary branches take the VB fate upon reducing both Wg/WNT and Dpp/BMP signaling.

In the artificially-induced VB default state of *btl-Gal4>Axn-GFP tkv* mutants, loss of the distal branches was detected much more severely in Tr3/9 than in other metameres (Figure 4G). This suggests that more cells in Tr3/9 are exposed to apoptosis. Indeed, strong expression of *rpr-lacZ* was variably detected in the whole distal part of Tr3/9 in these situations, when apoptosis was suppressed by H99 deficiency (Figure 4H).

Taken together, these results suggest that the VB identity is the default character of the primary branches and that extrinsic signals of Wg/WNT and Dpp/BMP induce the more derived primary branch types. In Tr3/9, the VB cells are apoptotically pruned whereas the derived primary branches are protected from apoptotic pruning (Figure 4I).

### The metamere identities for VB3/9 pruning are conferred by *Ubx* and *AbdB*

The *Drosophila* airways originate from the ectoderm, where differential expression of Hox genes confers the para-segmental identities. Thus, the 3 BX-C Hox genes present good candidates controlling the metameric specificity of VB pruning (Maeda and Karch 2006; Matsuda et al. 2015a).

At stage 11 when primary branching initiates, Ubx was detected from PS5 (Tr2) to the anterior half of PS13 (Tr10), its expression peaking in PS6 (Tr3) (Figure S5A) (Matsuda et al. 2015a). *abdA* expression is detected from PS7 (Tr4) to the anterior half of PS13 (Tr10), with highest expression in PS6 (Tr6) (Matsuda et al. 2015a). AbdB was expressed strongly in Tr10 (PS13) while it is weakly expressed in Tr9 at stage 11 (Figure S5B). By stage 13, AbdB becomes detected in the posterior metameres Tr7-Tr10, in a gradient with highest expression in Tr10 (Matsuda et al. 2015a). Taken together, the strong expression of Ubx in metamere 3 and the weak expression of AbdB in metamere 9 are good candidates for the metamere-identity inputs specifying VB3/9 pruning.

In *bxd^100^* mutants, the strong Ubx expression in PS6 (Tr3) becomes as weak as the one in PS5 (Tr2) (Camprodon and Castelligair 1994). Correspondingly, VB3 formation was restored (Figure 5A). This restored VB3 did not express the *rpr-lacZ* reporter (Figure 5A), suggesting that both *rpr* expression and VB3 pruning require high levels of Ubx expression in metamere 3.

**Figure 5.**
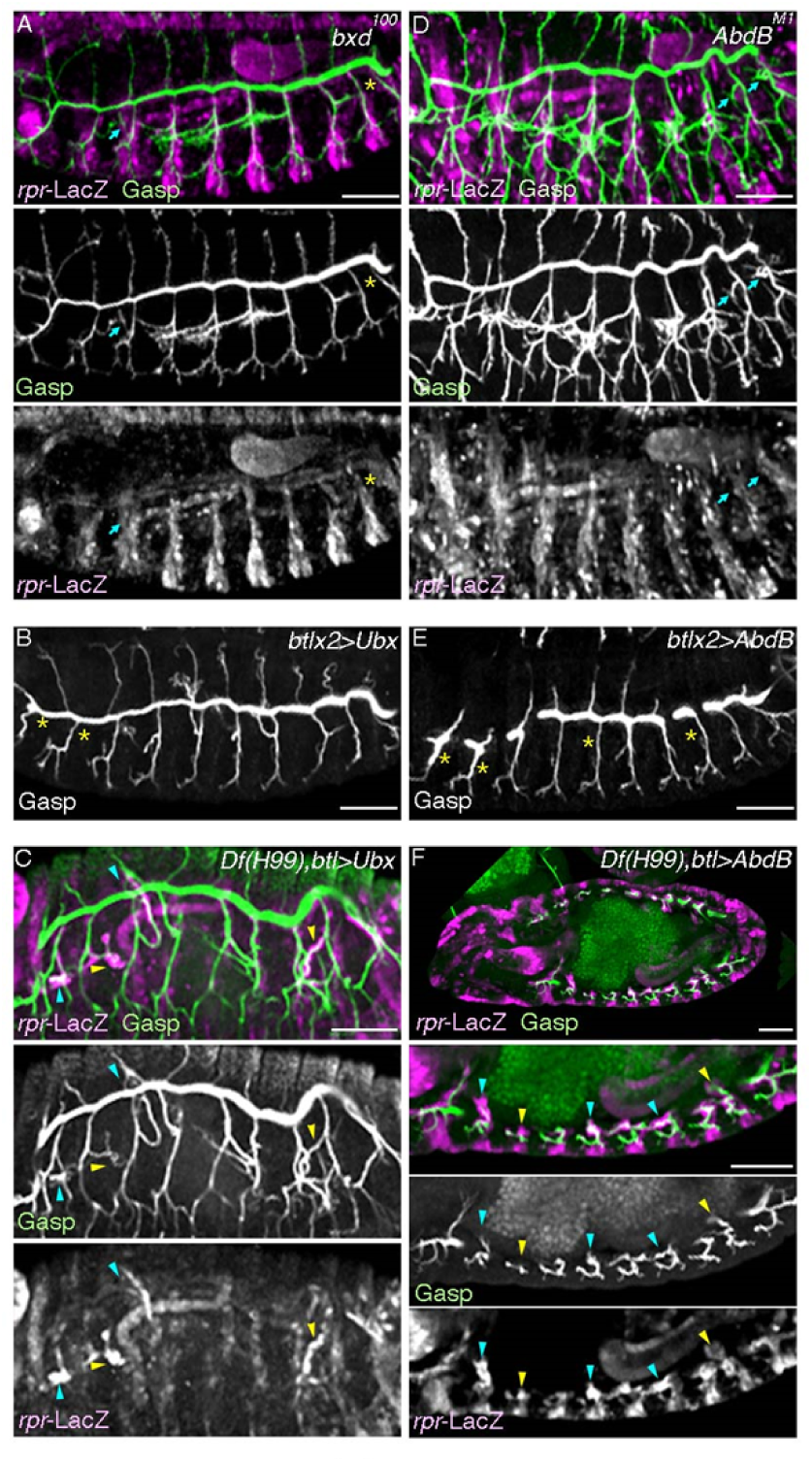
Ubx and AbdB instruct pruning of VB3/9. Gasp stained embryos showing involvement of *Ubx* (A-C) or *AbdB* (D-F) in VB pruning. (A-C) In *bxd^100^* mutants (A), VB3 is restored without causing detectable upregulation of *rpr-lacZ* (arrows) without affecting loss of VB9 (asterisks). The airway specific overexpression of Ubx (B) causes loss of VB2 and occasional loss of VB1 (asterisks). In *Df(H99)* mutant background (C), loss of VB1/2 upon Ubx overexpression is suppressed and ectopic *rpr-lacZ* signals in VB1/2 is detected (blue arrowheads). *rpr-lacZ* expression in VB3/9 is marked with yellow arrowheads. Note that the weak VB1 signal is out of the focus and a weak ectopic induction of *rpr-lacZ* is evident in VB4 (blue arrowheads). (D-F) In *AbdB^M1^* mutants (D), both VB9 and VB10 form (arrows) but they are negative for *rpr-lacZ*. The airway specific overexpression of AbdB (E) causes loss of VB in more anterior metameres (asterisks). In *Df(H99)* mutant background (F), AbdB overexpression accompanies *rpr-lacZ* upregulation in VB of more anterior metameres (blue arrowheads). *rpr-lacZ* expression in VB3/9 is marked with yellow arrowheads. Scale bars, 50 μm.

Conversely, upon *btl-*Gal4 mediated overexpression of Ubx in an otherwise wild type background, we detected a frequent loss of VB2 and a more occasional lack of VB1 (Figure 5B). Airway specific Ubx overexpression in the *H99* background, where apoptosis does not occur restored VB1/2 but ectopic upregulation of *rpr-lacZ* was detected in VB2 and more variably in VB1 (Figure 5C). Ectopic expression of *Ubx* is thus sufficient to induce *rpr* expression and VB pruning in the more anterior metameres, Tr1/2.

Loss or gain of function manipulations of Ubx did not majorly affect VBs of more posterior metameres, in accord with the notion that posteriorly expressed Hox genes tend to have dominant effects over the more anteriorly expressed Hox genes (Maeda and Karch 2006). However, VB still formed (Figure S5C) in *abdA* mutants, where the posterior Hox gene in Tr4-6 is mutated and Ubx expression is expected to become high. This suggests additional unknown modes of regulation on VB pruning. On the other hand, in AbdB mutants, VB-like extensions were detected not only in Tr9 but also in Tr10 (Figure 5D, S5D), whereas VB3 pruning was not affected (Figure 5D). In contrast to the wild type VB9 branches, which express *rpr* and undergo apoptosis, the persistent VB9 of *AbdB^M1^* mutants showed no up-regulation of *rpr-lacZ* (Figure 5D). These results suggest that *AbdB* is necessary for *rpr* induction and subsequent pruning in VB9. *AbdB* is also necessary to repress VB formation in Tr10.

Conversely, overexpression of AbdB with *btl-*Gal4 inhibited VB formation in the more anterior metameres (Figure 5E). AbdB overexpression in the *H99* background accompanied stronger *rpr-lacZ* expression in VBs (Figure 5F). We additionally note that AbdB overexpression variously suppressed extension and fusion of all the primary branches (Figure 5E-F) (Matsuda et al. 2015a). Taken together, AbdB expression is sufficient for induction of *rpr* expression and subsequent VB pruning.

We then examined if a pair of Hox cofactor TFs Exd and Hth function for VB3/9 pruning. *exd* zygotic mutants restored only VB3 (Figure S5E) whereas maternal and zygotic mutants of exd restored both VB3 and 9 (Figure S5F). In *hth* zygotic mutants, both VB3 and 9 are restored (Figure S5G-H). The restored VB3 and 9 in *hth* mutants was not accompanied by any *rpr-lacZ* up-regulation (Figure S5H). This suggests that the Exd and Hth co-factors contribute to the branch pruning activities of Ubx and AbdB (Figure S5I).

Taken together, we conclude that the metamere-identity inputs are composed of the Hox code of the Bithorax Complex. When Ubx or AbdB exceed a threshold level in Tr3 or in Tr9, respectively, they are expected to interfere with the VB identity, which autonomously induces *rpr* expression in VB and their subsequent apoptotic pruning. If VB pruning occurs solely through *ems* regulation, we speculate that interference with VB differentiation programs may lead to *rpr-lacZ* induction. Alternatively, *Ubx* and *AbdB* may control other targets whose functions can be compensated by *ems* overexpression. A candidate such target is *E5*.

### Branch extension and pruning regulated by the distalizing factor *pointed*

*rpr* expression in VB3/9 itself represents a primary branch identity. Because expression of *hairy*-LacZ marking DB/DT/VB is promoted by the distalizing factors *rho* and *pointed* (*pnt*) (Matsuda et al. 2015b), we examined potential roles of the distalizing factors on VB pruning.

*pnt* is induced in the airway primordia by Rho mediated EGFR signaling (Matsuda et al. 2015b) whereas Btl/FGFR also induces *pnt* expression around the tips of the primary branches (Samakovlis et al. 1996a) (Figure 6H). Branching defects of *pnt* mutants are more severe than those of *rho* single mutants but are milder than those of *btl/FGFR* single mutants (Figure 6A-D) (Klambt et al. 1992; Klambt 1993; Wappner et al. 1997; Llimargas and Casanova 1999; Bradley and Andrew 2001; Matsuda et al. 2015b). However, double mutants of either *pnt* and *btl* or *pnt* and *rho* showed synergistic enhancement of the branching defects, resembling close the *rho btl/FGFR* double mutants (Figure 6E-H). Therefore, *pnt* is an essential mediator of Rho mediated EGFR signaling and Btl/FGFR signaling, both promoting primary branch formation.

**Figure 6.**
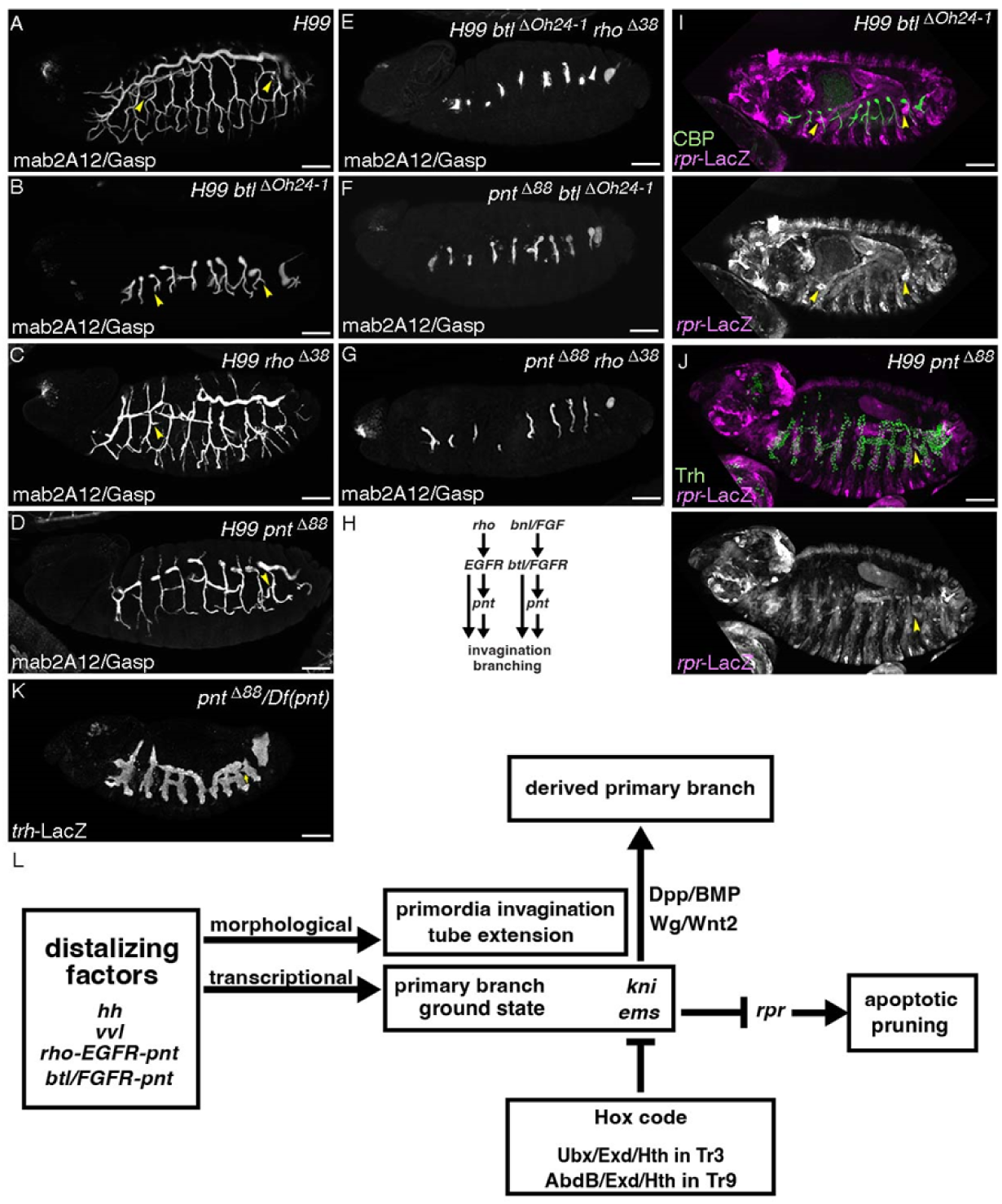
The r*h*o*-EGFR-pnt* distalizing module promotes both extension and pruning of the primary branches. (A-H), VB3/9 restoration in *H99* mutants (A) is not suppressed by *btl/FGFR* mutation (B), but is partially suppressed by mutations of *rho* (C) or *pnt* (D). *pnt* mutation enhances the branching phenotypes of *rho* or *btl/FGFR* single mutants; *pnt btl/FGFR* (F) or *pnt rho* (G) double mutants have simpler tubes with less ramification like those of *H99 rho btl/FGFR* mutants (E, H). (I-J) In *H99 btl* double mutants, *rpr-lacZ* is strongly upregulated in the restored VB3/9 (I, arrowheads) whereas the restored VB9 in the *H99 pnt* double mutant embryo does not upregulate *rpr-lacZ* (J, arrowheads). (K) *pnt* mutants infrequently restore VB9 (arrowhead). (L) A regulatory model of vessel extension and pruning in the developing *Drosophila* airways. See text for details. Scale bars, 50 μm.

In *H99 btl* mutants, VB3/9 were formed (Figure 6B) and *rpr*-LacZ was induced there (Figure 6I). In contrast, *rho* mutation affects VB3/9 formation of *H99* mutants, suggesting that VB3/9 budding requires Rho more than Btl/FGFR. Despite that *pnt* is stronger involved in primary branching compared to *rho*, VB3/9 formation was more often detected in *H99 pnt* mutants than in *H99 rho* mutants (Figure 6C-D) (Matsuda et al. 2015b). In those restored VBs, *rpr*-LacZ expression was very much reduced (Figure 6J). Consistent with the reduction of *rpr* induction in VBs, we detected VB9 restoration in *pnt* mutants at a lower frequency (Figure 6K).

Taken together, our results suggest that Rho-EGFR-Pnt signaling promotes both primary branch formation and pruning (Figure 6L). We speculate that *rho* mutation synergizes with the Hox code to dampen VB3/9 budding even in the *H99* background, whereas *pnt* mutation leaves Rho-EGFR signaling high enough for VB3/9 budding. However, *pnt* mutation prevents the primary branch identity from being properly established, resulting in reduced *rpr* induction. The resultant failure to robustly induce *rpr* expression infrequently allows VB9 restoration.

### The same overlying regulators control tube outgrowth and pruning

In the developing *Drosophila* airways, the interplay of several signaling systems along the anterior-posterior and the dorso-ventral body axes set the airway progenitors on the 2D fields, where the distalizing factors primes the progenitors for distal gene expression and for radial invagination to generate tube outgrowth into the body (Matsuda et al.

2015b; Matsuda et al. 2026). In the absence of the distalizing factors, the progenitors rarely invaginate (*hh vvl* double mutants), stay at the invaginated places (*rho btl/FGFR* double mutants) or form rudiments of primary ramification of the distal tips into 3 directions (*btl/FGFR* mutants) (Figure 2C). Irrespective of the directions of branch extension, the tips of the distal airways or the primary branches share the default VB state that simultaneously express Kni and Ems, meaning that *kni* expression under the VB enhancer is the default marker for the tips of the distal airways (the primary branches).

Our results suggest that lack of VB3/9 does not require novel extrinsic signals that suppress tube outgrowth. Rather, interference with the tube outgrowth programs can induce apoptotic branch pruning. The distalizing module of Rho-EGFR-Pnt promotes tube formation. At the same time, the primary branch identity specified by this module is a prerequisite for *rpr* induction in VB3/9 and branch pruning. Specifically, establishing the VB identity is prerequisite for VB formation and its subsequent pruning. Extrinsic guidance molecules like Wg/Wnt2 or Dpp/BMP that diversify the primary branch identities protect cells from VB identity-dependent apoptosis. The Hox code impinges on the branch identity hierarchy to destabilize a specific primary branch identity, and thereby induces apoptosis and pruning in specific metameres.

It is assumed that during evolution, modification of parts of appendages to generate inwardly extending tracheal tubes had aided insect terrestrialization (Franch-Marro et al. 2006) and that fusion of those ancient tracheal branches is a relatively new invention (Caviglia and Luschnig 2013). Intriguingly, VBs are specialized for producing tip cells of only the terminal cell type lacking the ability of branch fusion. This suggests that VBs resemble the ancient form of the insect airways. Indeed, the cells of the primary branches that migrate dorsally or ventrally just beneath the overlying ectoderm additionally receive guidance molecules like Wg/Wnt2 or Dpp/BMP, which induce more derived primary branch identities whereas the interiorly moving primary branches do not encounter such signals and keep the default VB state. Repurposing the branch outgrowth programs to branch pruning may contribute to the intricate balance of branch extension and removal during branching morphogenesis of more complex tubular organs in vertebrates.

## Materials and Methods

### Fly genetics

Flies kept over balancer chromosomes (Lindsley et al. 1992) were grown in standard medium. We obtained the appropriate genotypes by standard genetic crosses. For overexpression of genes, we used the Gal4/UAS system (Brand and Perrimon 1993). Mutant embryos were identified by the expression of *twi-lacZ, ftz-lacZ, Ubx-lacZ or Dfd-GFP* constructs inserted on balancer chromosomes. See Flybase for details of strains described below.

*1-eve-1*=*trh-lacZ* (a gift from Dr. N. Perrimon) (Perrimon et al. 1991)

*abdA^M1^* (a gift from Dr. B. Gebelein) (Li-Kroeger et al. 2008)

*AbdB^M1^*(a gift from Dr. I. Lohmann) (Lohmann et al. 2002)

*AbdB^M2^* (a gift from Dr. I. Lohmann) (Lohmann et al. 2002)

*arm^4^* (BDSC)

*btl^ΔOh10^* (Ohshiro and Saigo 1997)

*btl^ΔOh24-1^* (Ohshiro and Saigo 1997)

*btl^724^*(a gift from Dr. A. Ghabrial) (Ghabrial et al. 2011)

*btl*-CD8GFP (a gift from Dr. M. Sato)

*btl-gal4* (a gift from Dr. S. Hayashi) (Shiga et al. 1996)

*Df(3L)BSC448= Df(kni/knrl)* (BDSC)

*Df(3R)Exel9012*=*Df(pnt)* (BDSC)

*Df(3L)riXT*=*Df(kni/knrl)* (a gift from Dr. R. Schuh) (Chen et al. 1998)

*Df(5)*=*Df(salm/salr)* (a gift from Dr. M. Llimargas) (Barrio et al. 1999; Llimargas 2000)

*exd^1^* FRT (BDSC)

*hth^P2^* (a gift from Dr. R. Mann and Dr. B. Gebelein) (Gebelein et al. 2002)

*kni-(dpp)-lacZ* (a gift from Dr. R. Schuh) (Chen et al. 1998)

*kni early rescue fragment; Df(3L)riXT* (a gift from Dr. R. Schuh) (Chen et al. 1998)

*ovoD1 FRT; hsFlp* (BDSC)

*P0144-lacZ* (a gift from Dr. W. Janning, Flyview)

*pnt^Δ88^* (BDSC)

*rpr^87^* (a gift from Dr. A. Bergmann and Dr. K. White) (Fan et al. 2010; Tan et al. 2011)

*rpr-11kb-lacZ* (a gift from Dr. I. Lohmann) (Lohmann 2003)

*salm^1^* (BDSC)

*salm-TSE-lacZ* (a gift from Dr. R. Schuh) (Kuhnlein and Schuh 1996)

*tkv^7^* (BDSC)

*tkv^8^* (BDSC)

*UAS-AbdB* (BDSC)

*UAS-Axn-GFP* (BDSC) *UAS-dpp* (BDSC)

*UAS-ems* (a gift from Dr. H. Reichert) (Hartmann et al. 2010)

*UAS-kni* (a gift from Dr. R. Schuh) (Chen et al. 1998)

*UAS-knrl* (a gift from Dr. R. Schuh) (Chen et al. 1998)*UAS-p35* (BDSC)

*UAS-salm* (BDSC)

*UAS-tau-lacZ* (a gift from J. B. Thomas) (Callahan et al. 1995)

*UAS-Ubx* (BDSC)

*UAS-wg* (DGRC Kyoto)

In situ hybridization and immunostaining

Egg collection was done with apple/grape juice plate at 25°C. Embryos were bleached and fixed as previously described (Patel 1994) for 15-30 minutes with a 1:1 mixture of heptane and a fix solution (3.7 % formaldehyde, 0.1M Hepes pH6.9, 2mM MgSO4). Embryos were dechorionated with methanol and incubated in 0.1% PBT supplemented with 0.5% BSA. Staging of embryos was done as previously described (Campos-Ortega and Hartenstein 1997).

For immunostaining the following primary antibodies were used: mouse anti-Abd-B (1:10, DSHB, donated by Dr. S. Celniker) (Celniker et al. 1989)

mouse anti-DSRF (1:1000, a gift from Dr. Michael Gilman)

rabbit anti-Dys (1:500, a gift from Dr. L. Jiang) (Jiang and Crews 2003)

rabbit anti-Ems (a gift from Dr. U. Walldorf) (Walldorf and Gehring 1992)

mouse anti-Fas3 (1:10, DSHB, donated by Dr. C. Goodman) (Patel et al. 1987)

Guinea-pig anti-Gasp (1:1000) (Tiklova et al. 2013)

Guinea-pig anti-Kni (1:300) (developed by Dr. J. Reinitz and distributed by Dr. Y. Hiromi, East Asian Segmentation Antibody Center, Mishima, Japan) (Kosman et al. 1998)

mouse mab2A12 (anti-Gasp) (1:5, DSHB, donated by Drs. M. Krasnow, N. Patel and C. Goodman) (Samakovlis et al. 1996b; Tiklova et al. 2013)

rabbit anti-Salm (1:200, gifts from Dr. R. Barrio and Dr. T. Cook) (Barrio et al. 1999; Xie et al. 2007)

rat anti-Trh (1:200, a gift from Dr. D. Andrew) (Henderson et al. 1999) rabbit anti-Trh (1:50) (Matsuda et al. 2026)

mouse anti-Ubx (1:10, a gift from Dr. R. White) (White and Wilcox 1984)

Chicken anti-LacZ (1:500, Abcam)

Goat anti-LacZ (1:500, Biogenesis)

Rabbit anti-LacZ (1:1000, Capel)

Rabbit anti-GFP (1:500, JL-8 Clontech)

Donkey or goat biotin- or fluorescently labeled secondary antibodies made against the host species of primary antibodies were purchased from Jackson Laboratories. Streptavidin coupled with AMCA, FITC or Cy5 were used when necessary. For mab2A12 detection TSA amplification (PerkinElmer) was used.

Double fluorescent labeling with RNA probe and antibody was carried out as described (Goto and Hayashi 1997). The following cDNA clones were used to make hybridization probes;.

*bnl* (a gift from Dr. M. Krasnow) (Hosono et al. 2003)

*dpp* (Hosono et al. 2003)

*hairy* (a gift from D. Ish-Horowicz) (Hooper et al. 1989)

*rpr* (a gift from Dr. H. Steller) (White et al. 1994)

*wg* (Hosono et al. 2003)

Confocal images were taken by Biorad MRC1024, Oympus Fluoview 1000 or Zeiss LSM780/LSM800. Images were processed by ImageJ and figures were prepared with Photoshop and Illustrator.

## Acknowledgments

We thank the members of the fly community who isolated, characterized or distributed fly strains, antibodies or DNA clones. Especially, we thank Drs. D. Andrew, R. Barrio, A. Bergmann, T. Cook, A. Ghabrial, B. Gebelein, S. Hayashi, Y. Hiromi, D. Ish-Horowicz, W. Janning, L. Jiang, D. P. Kiehart, M. Krasnow, M. Llimargas, I. Lohmann, R. Mann, N. Perrimon, H. Reichert, M. Sato, R. Schuh, H. Steller, J. B. Thomas, U. Walldorf, K. White, R. White, BDSC, DGRC, DSHB for directly sharing fly strains, antibodies or DNA clones. We thank Flybase for the *Drosophila* genomic resources. We thank the Stockholm University Imaging Facility and the MBW technical group for fly service and V. Tsarouhas for microscope help. Special thanks to Y. Emori and F. Ui-Tei for help in maintaining fly strains after the retirement of K.S.

This work was funded by the Ministry of Education, Culture, Sport, Science and Technology of Japan to K.S. and the Swedish Research Council, the Swedish Cancer Society and German Research Foundation to C.S.

## Author contribution

RM, conceived the project based on the phenotype of *exd* zygotic mutants, designed the experiments, performed experiments, interpreted data, drafted the manuscript with inputs from CH, writing, figure preparation

CH, found VB3 in *exd* zygotic mutants, performed experiments, figure preparation, writing

KS, funding acquisition, provided experiment and analysis tools

CS, funding acquisition, provided experiment and analysis tools, writing

---

**Figure S1.**
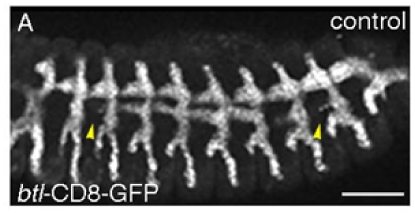
Cell debris positive for *btl*-CD8GFP is detected around VB3/9 at stage 13. Scale bar, 50μm.

**Figure S2.**
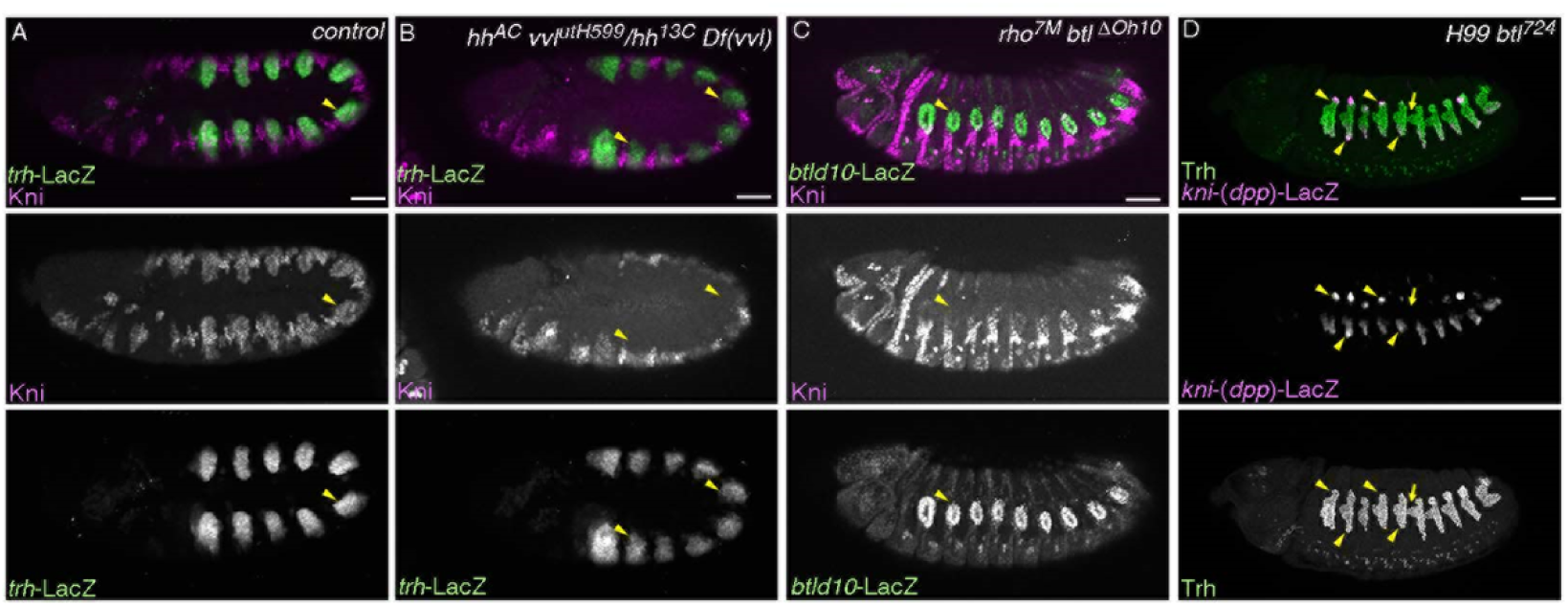
The distalizing factors promote *kni* expression in the distal tips of the distal airways. Compared to the control, Kni expression is abolished in *hh vvl* double mutants by stage 11 (A-B, arrowheads). Kni expression is variably reduced in *rho btl/FGFR* double mutants at stage 13 (C, arrowheads). In *H99 btl* mutants where apoptosis is suppressed by *H99* deficiency, 3 primary branch structures (DT, VB and GB) are apparent. Cells positive for *kni-(dpp)*-LacZ are detected in the dorsal tip of the DT (yellow arrowheads) and in GB cells (yellow arrowheads). VB is marked with yellow arrows. Scale bars, 50μm.

**Figure S3.**
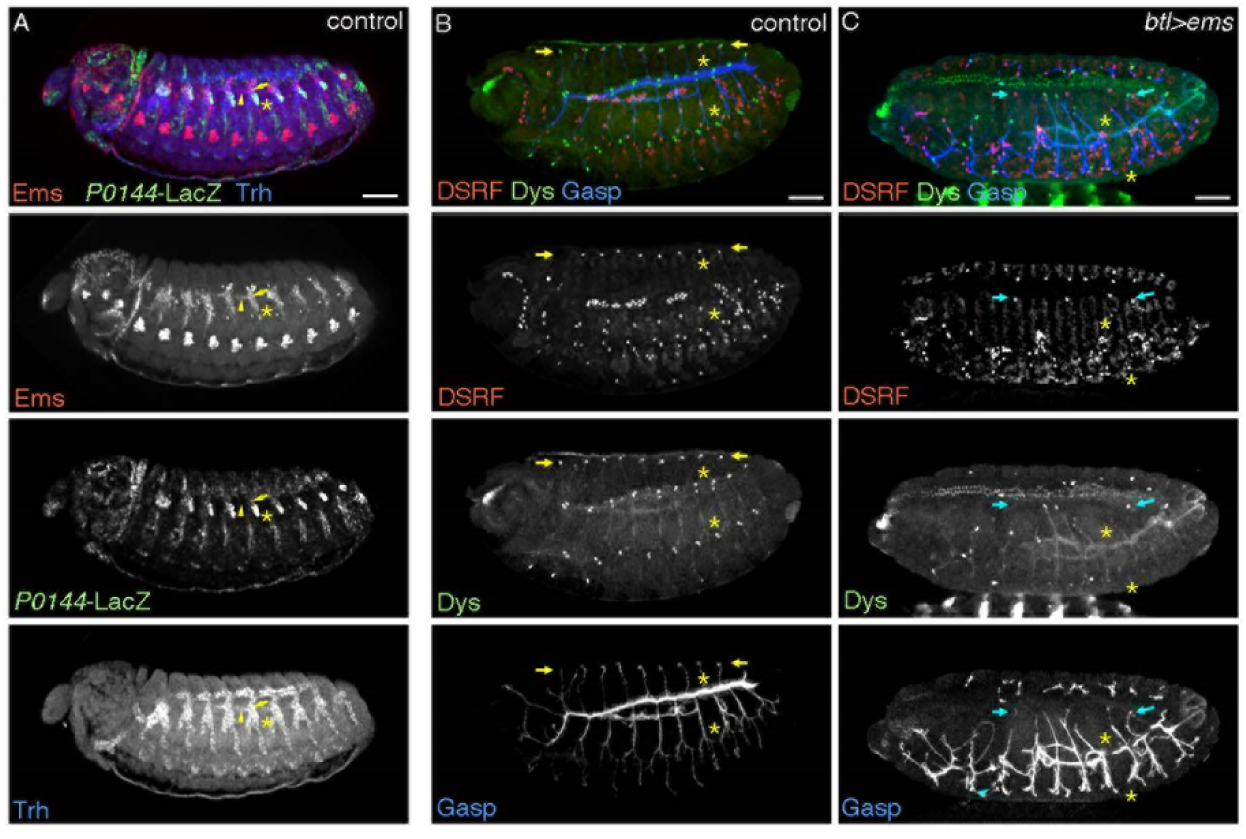
Ems expression and its effect on tip cell identities. (A) Ems is expressed not only in VB (arrowheads) and TC (arrows), but also in the *P0144-lacZ* positive proximal SB cells (asterisks). (B-C) Compared to the control (B), Ems overexpression (C) variably suppresses branch fusion and expression of the fusion fate marker Dys (asterisks), with some effects on expression of the terminal marker DSRF (arrows). Scale bars, 50 μm.

**Figure S4.**
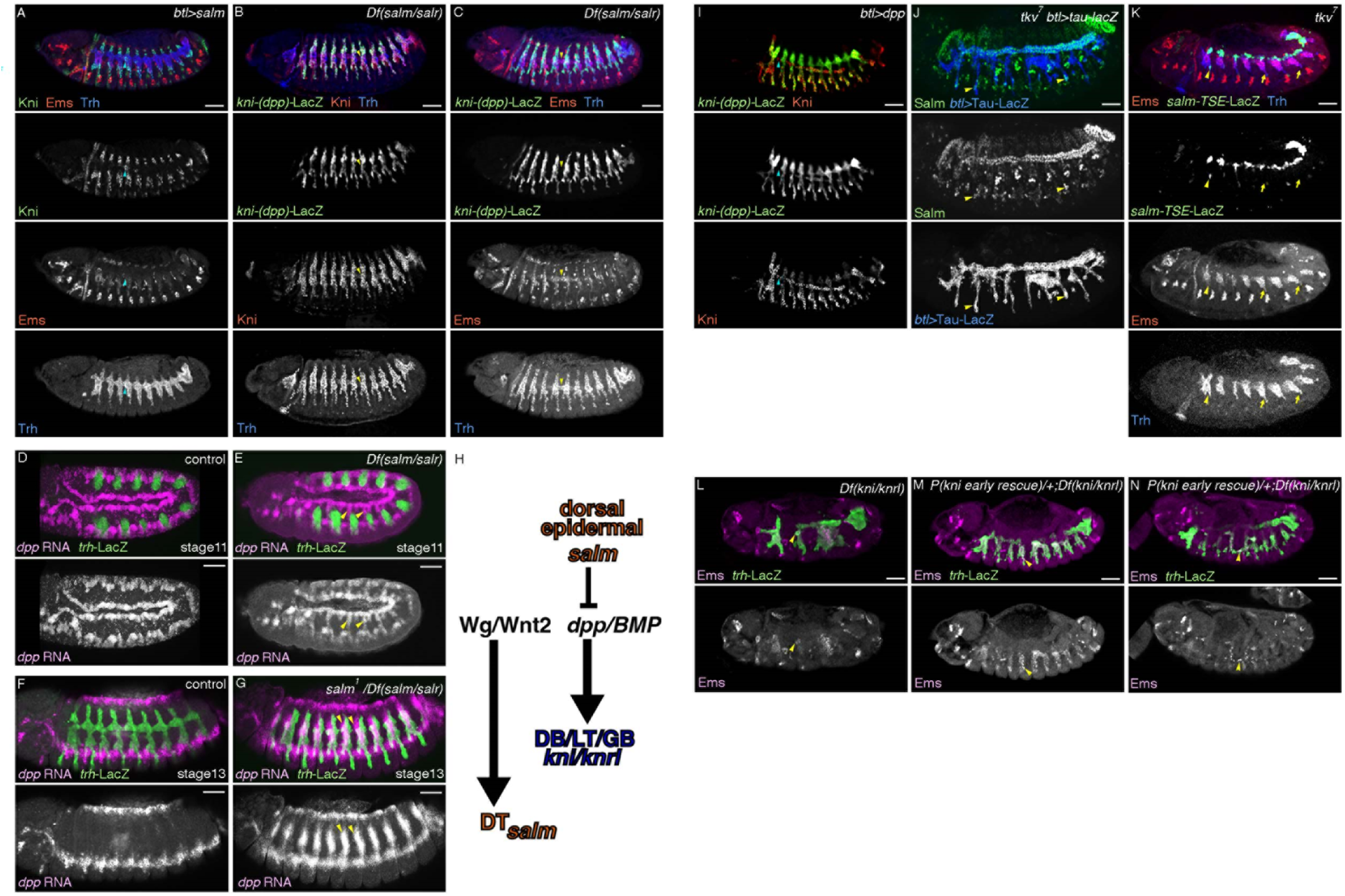
Involvement of Salm and Kni/Knrl in VB identity control. (A-H) *salm* analysis. Upon overexpression of Salm (A), expression of both Ems and Kni is reduced in VB (arrowheads). In *Df(salm/salr)* embryos (B, C), expression of both Kni and *kni-(dpp)-lacZ* covers all the distal part with exclusion of *kni-(dpp)-lacZ* expression from the VB (arrowheads in B, C) where Ems is expressed (C). Compared to the control (D, F), expression of *dpp/BMP* is ectopically detected from stage 11 in the lateral ectoderm in *salm* mutants (E, G, arrows), which is summarized in (H). (I-M) Analysis of the *dpp/BMP-kni/knrl* pathway. Upon Dpp overexpression (I), all distal cells become positive for Kni and *kni-(dpp)-lacZ*, although the VB cells are more resistant to ectopic *kni-(dpp)-lacZ* induction (arrowheads). In tkv mutants (J, K), expression of Salm (J) or *salm-TSE-lacZ* (K) becomes ectopically detected at ventral branches (arrowheads). Ems expression also variably expands to the ventral branches (K, arrows). In *Df(kni/knrl)* mutants (L), the residual VB cells are positive for Ems (arrowheads). In *Df(kni/knrl)* mutants with a *kni* segmentation-rescue fragment (M,N), ventral branches variably become positive for Ems (arrowheads). Scale bars, 50 μm.

**Figure S5.**
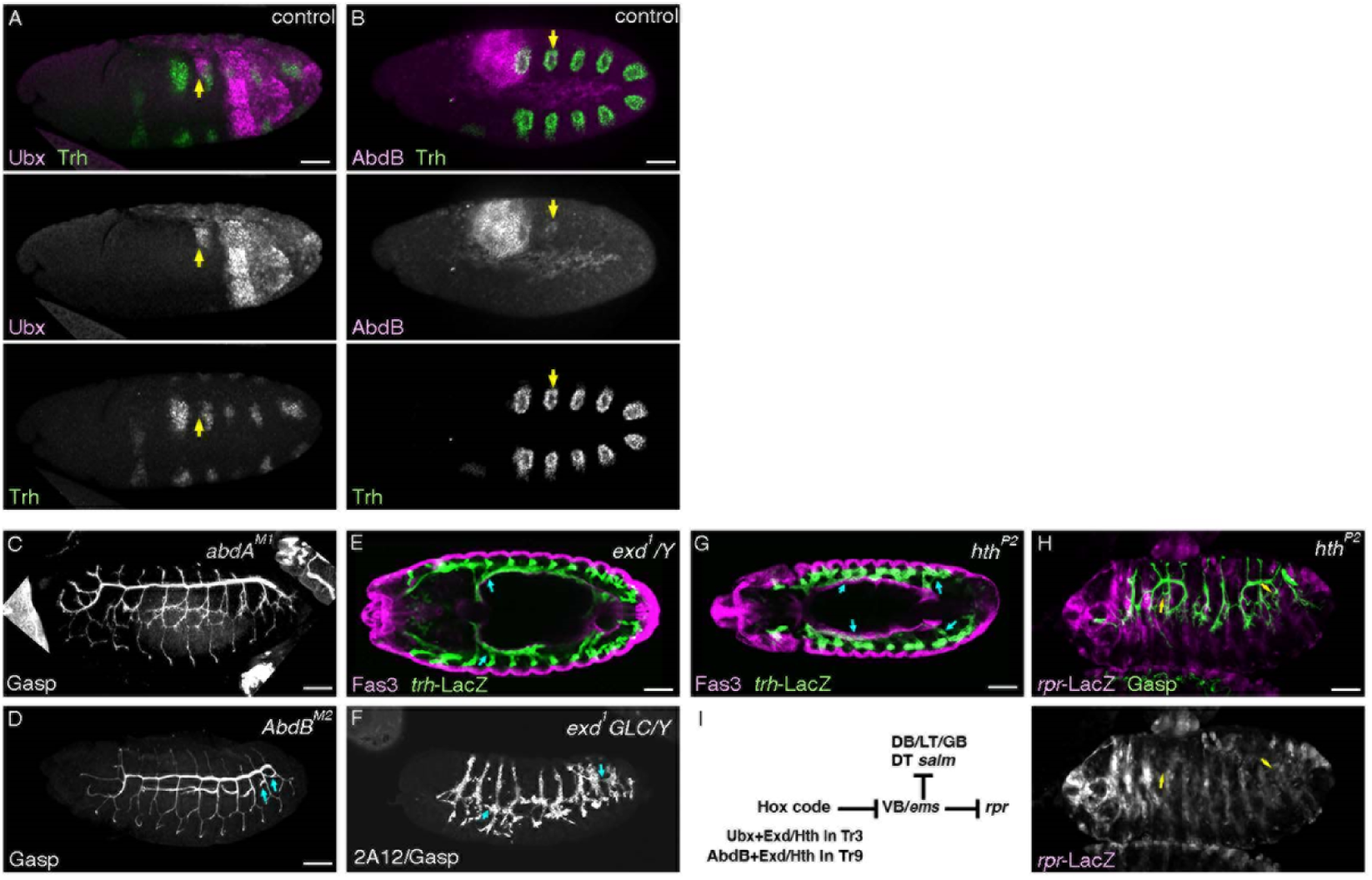
Expression patterns of BX-C Hox genes and phenotypes of BX-C Hox genes and their cofactors *exd* and *hth*. (A,B) Expression of Ubx and AbdB. Ubx expression (A) in Tr2 (arrows) is lower than that in Tr3 while AbdB expression (B) in Tr9 (arrows) is lower than that in Tr10. (C-I) Phenotypes of BX-C mutants, *exd* and *hth*. VB formation in *abdA^M1^* mutants (C) is comparable to the control whereas in *AbdB^M2^* mutants, both VB9 and VB10 form (D, arrows). In *exd* zygotic mutants (E), an ectopic VB3 forms (arrows) while in maternal and zygotic *exd* mutants (F) or in zygotic *hth* mutants (G), both ectopic VB3 and VB9 form (arrows). The ectopic VB3 and VB9 in *hth* mutants do not express *rpr-lacZ* (H). A model of Hox dependent regulation of the VB3/9 pruning is shown in (I). Scale bars, 50 μm.

## Notes

### Competing Interest Statement

The authors have declared no competing interest.

